# Appendometer: A system for simultaneous, high-throughput morphometry of *Drosophila* legs and wings

**DOI:** 10.1101/2025.01.21.634122

**Authors:** Daniela M. Rossoni, Connor Murray, Arthur Porto, David Houle

## Abstract

1. The inheritance of many different organismal features is correlated, as is their evolution, suggesting that we need to understand the pattern and causes of those correlations to understand evolution. Unfortunately, we generally lack the ability to rapidly and accurately measure large numbers of traits, making it difficult to describe the patterns of trait relationships or formulate hypothesis about the causes of their entanglement. We have previously developed a system to make high-dimensional measurements of Drosophilid fly wings in live specimens. Here, we report the extension of this approach to rapidly assess the dimensions of the distal leg segments, femur, tibia, and tarsi. Using the system, we describe the covariance of the wing and leg morphology and evaluate the relative rates of evolution of legs and wings.
2. We use two simple suction devices to immobilize and position legs and wings of an anaesthetized fly for imaging, then take a single image incorporating both appendages. We employed a machine learning method to measure leg segment lengths, which should be broadly applicable across diverse taxa. Experienced users can image the legs and wings of a fly every two minutes, with outlier detection and correction taking approximately 40 seconds. To demonstrate the usefulness of these methods, we measured the legs and wings of over 4,000 specimens from 43 different Drosophilid taxa. We estimated the rate of wing and leg evolution using a phylogenetic mixed model.
3. Repeatabilities of leg segments lengths averaged over 80%. The rate of evolution of wing and leg sizes are similar, but the rate of wing proportion evolution is 1.6 times as high as that of leg proportions due to strong allometric changes in wing shape. Within-species variation in leg proportions is highly correlated with the rate of leg proportion evolution, as is true for wings. Relative lengths of leg segments showed a strong pattern of negative correlations between the lengths of the tarsal segments and of the femurs and tibias, while all other segment correlations were positive. This pattern was repeated in the rate estimates, suggesting that selection favors tradeoffs between tarsi and the remainder of the leg.
4. Our simple system for imaging and measuring legs and wings simultaneously has high throughput and repeatability. It is readily applicable to a wide variety of winged insects and other traits, including wings, that could be imaged. Applied to *Drosophila*, our morphometric system enlarges the ability to study inheritance, pleiotropy, and evolution in this important model taxon.

## 1 INTRODUCTION

It has long been appreciated that the environmental responses, inheritance and evolution of different organismal traits are frequently correlated [1], resulting in an integrated phenotype. Some of the most virulent controversies in recent evolutionary biology concern the potential causes of these patterns (e.g., [2]). Two major, non-exclusive explanations exist for the entanglement of traits across multiple biological levels: pleiotropy, the tendency for one genetic change to affect multiple traits [3], and the possibility that natural selection favors particular combinations of phenotypic traits. Studying the integration of variation and evolution is facilitated by data on a wide range of traits. Unfortunately, our datasets often measure only a limited number of traits, rather than being comprehensive. Consequently, while we have enough data to be sure that biological systems are chock-a-block with cases of correlated variation and evolution, we struggle to document how pervasive those patterns are.

A limiting factor in several empirical studies of such problems is the ability to rapidly measure the diversity of traits that can affect organismal fitness [4, 5]. A typical study measures just a few such traits, or a suite of related traits. While we would like to take the time to measure a wide array of traits, the costs of traditional phenotyping approaches are often high. Technical advances in recent years have enabled high-throughput and fine-grained measurement of some sets of traits, such as gene expression and protein abundance in a wide variety of organisms, although at substantial cost. In addition, taxon-specific techniques allow the assessment of other suites of traits, such as behavior in *Drosophila* flies and *Caenorhabditis* [6].

Houle et al. [7, 8] developed a system for rapid morphometry of wings of Drosophilid flies, the Wingmachine. This system is ideal for investigating wing-specific questions. For instance, it has been used to characterize the inheritance of variation in wings [7–10], to perform artificial selection experiments on multiple aspects of wings [11–17], and to characterize the pattern of evolution of wings in the family Drosophilidae [18–20].

Wing-specific data enables characterization of pleiotropy within the wing [10], but genetic variants with effects on the wing may have pleiotropic effects on other aspects of morphology and function. Furthermore, geometry dictates that changes to any part of the wing must also cause changes across the whole wing, because insect wings are continuous epithelial structures. This geometrical necessity is mediated by such processes as transport of gene products, and mechanical forces caused by growth and cell movement [21].

To move beyond the constraints of geometry, we have expanded our morphometric measurements to include legs as well as wings. Here, we report the construction of a high-throughput system, the Appendometer, that enables simultaneous measurement of both wings and legs of live flies. We know that many of the same developmental pathways and processes are involved in the development of both appendages [22–24]. On the other hand, wings and legs are physically and developmentally autonomous and have different functions, making within- and among-appendage evolution a suitable model for investigating the influence of pleiotropy and natural selection on evolution.

We developed a simple system for simultaneous imaging of legs and wings of live specimens, which should be readily applicable to a variety of winged insects. We complemented this with an automated approach to extracting leg length measurements using machine learning [25] that can be readily extended to any morphological traits captured in 2-D images. We demonstrate the usefulness of this system by estimating both phenotypic variation and the rate of evolution in legs and wings in the genus *Drosophila*.

## 2 MATERIALS AND METHODS

### 2.1 Specimen handling

To present wings and legs for imaging, we used a pair of the suction devices described in [11]. Each grabber forms a thin opening between of a glass microscope cover slip positioned above a glass slide by pieces of double-sided tape. Suction is applied to one end of the opening and used to draw appendages into the open end of the slot. For a wing grabber, we apply a single thickness of tape on each side of the imaging slot; for a leg grabber, two pieces of tape are applied, doubling the height of the slot.

The hardware used to position specimens for imaging is shown in Fig. 1. We used a positioning block milled from blue acetal copolymer to hold a pair of grabbers in the correct orientation. Fig. 1a-b shows the positioning block with two grabbers on the opposite sides of the fly. The positioning block and diffuser are placed on a plexiglass surface that extends into the imaging path of the macroscope, as shown in Fig. 1b. After anaesthetizing the fly using Fly Nap, the user first positions the left wing of the fly in the left grabber, then places the second grabber to the right of the fly and slides it to the right while coaxing the right set of legs into that grabber (Fig. 1b). The diffuser is then withdrawn, and the positioning block with the immobilized fly in the grabbers is moved to the imaging area on the left side of the plexiglass sheet, where the user takes an image (Fig. 1c).

**FIGURE 1.**
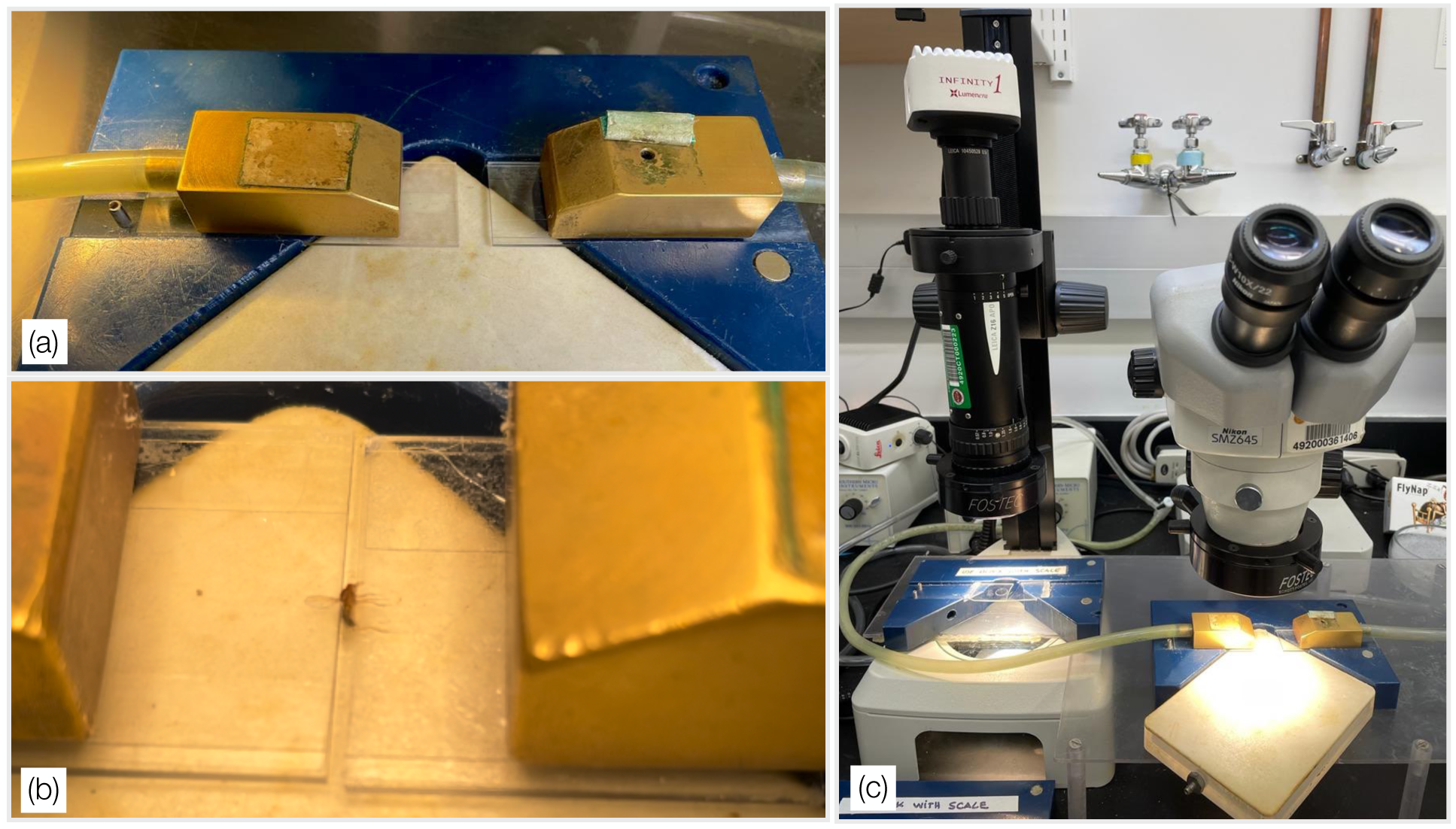
-Hardware used for simultaneous imaging of legs and wings. (a) Wing grabbers on the blue positioning block. The white area in the lower part of the image is the CO_2_ diffuser plate on which the specimen is manipulated. (b) Anesthetized fly in dorsal position with the left wing positioned into the left grabber, and the set of right legs placed into the right grabber. (c) Relationship between the fly-positioning area with the microscope at the right and the imaging area with the camera to the left.

Fig. 2 shows an example Appendometer image (a), and (b) the output of the machine learning algorithm. A detailed description of the imaging process is in Supplementary Material.

**FIGURE 2.**
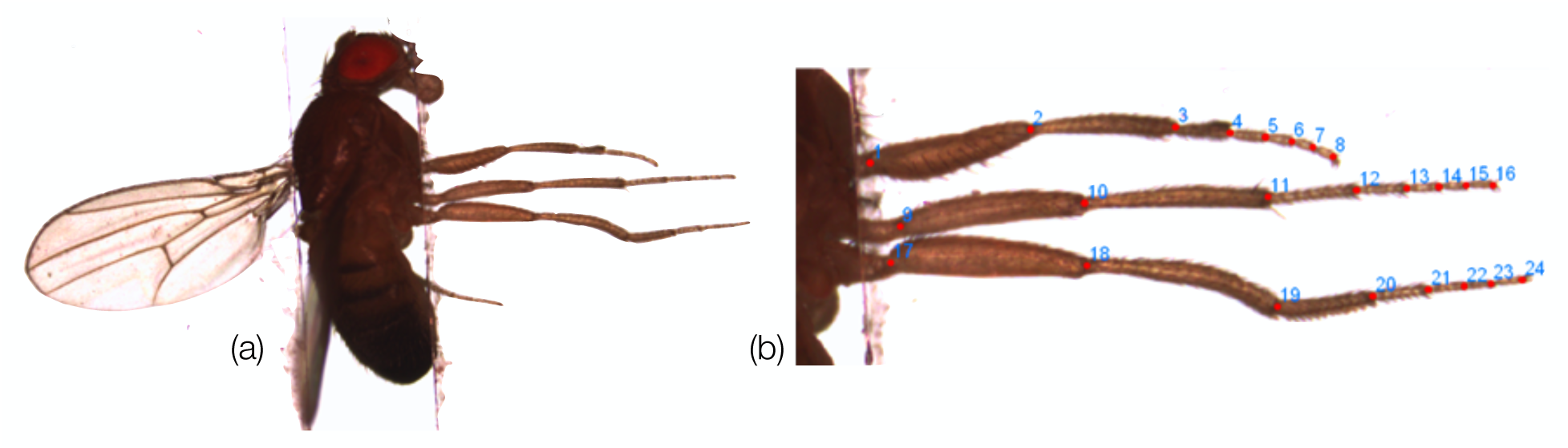
-(a) Appendometer image showing the left wing and right legs of an anesthetized fly. (b) Output of the machine-learning algorithm.

Once an image is saved, two guide landmarks are recorded at the base of the wing (as described in [11]), and one at the proximal end of the femur of each leg to facilitate automated identification of a region of interest containing all three legs. Leg guide points were placed along the suture between the trochanter and femur at the point furthest from the joint between the femur and tibia (Supporting Information Video 1). Imaging and recording of the scale and specimen information were automated using ImagePro 7.0 software (Media Cybernetics).

### 2.2 Imaging

We used two low-magnification macroscopes to image flies: an Optem Zoom 125, fitted with an Olympus Q-color 3 camera, and a Leica Z16 APO Macroscope, fitted with a Lumenera Infinity 1-3 camera (Fig. 1c, left). The Leica has higher resolution than the Optem. Images were recorded at a resolution of 2048×1536 pixels and saved as TIFF format files.

### 2.3 Hand-landmarking

To evaluate human error in the placement of landmarks, one observer hand-landmarked the end points of femurs, tibias, and tarsomeres on all three legs on 200 images (Fig. 3) using tpsDig2 [26]. Each image was landmarked twice, with at least two days between the sessions. To include a range of image qualities recorded by many imagers, we sampled images from three different experiments. Two sets of 50 images were sampled from experimental blocks from a large quantitative genetic experiment in *Drosophila melanogaster*, imaged 18 months apart. The other two sets were 50 images of *Drosophila acutilabella* and of *Scaptomyza latifasciaeformis* analyzed in this contribution. Femur ends were placed as described for guide points. Leg ends were placed at the base of the terminal claws on the fifth tarsomere. All other points were placed at the midpoint of the connection between neighboring segments (Fig. 3).

**FIGURE 3.**
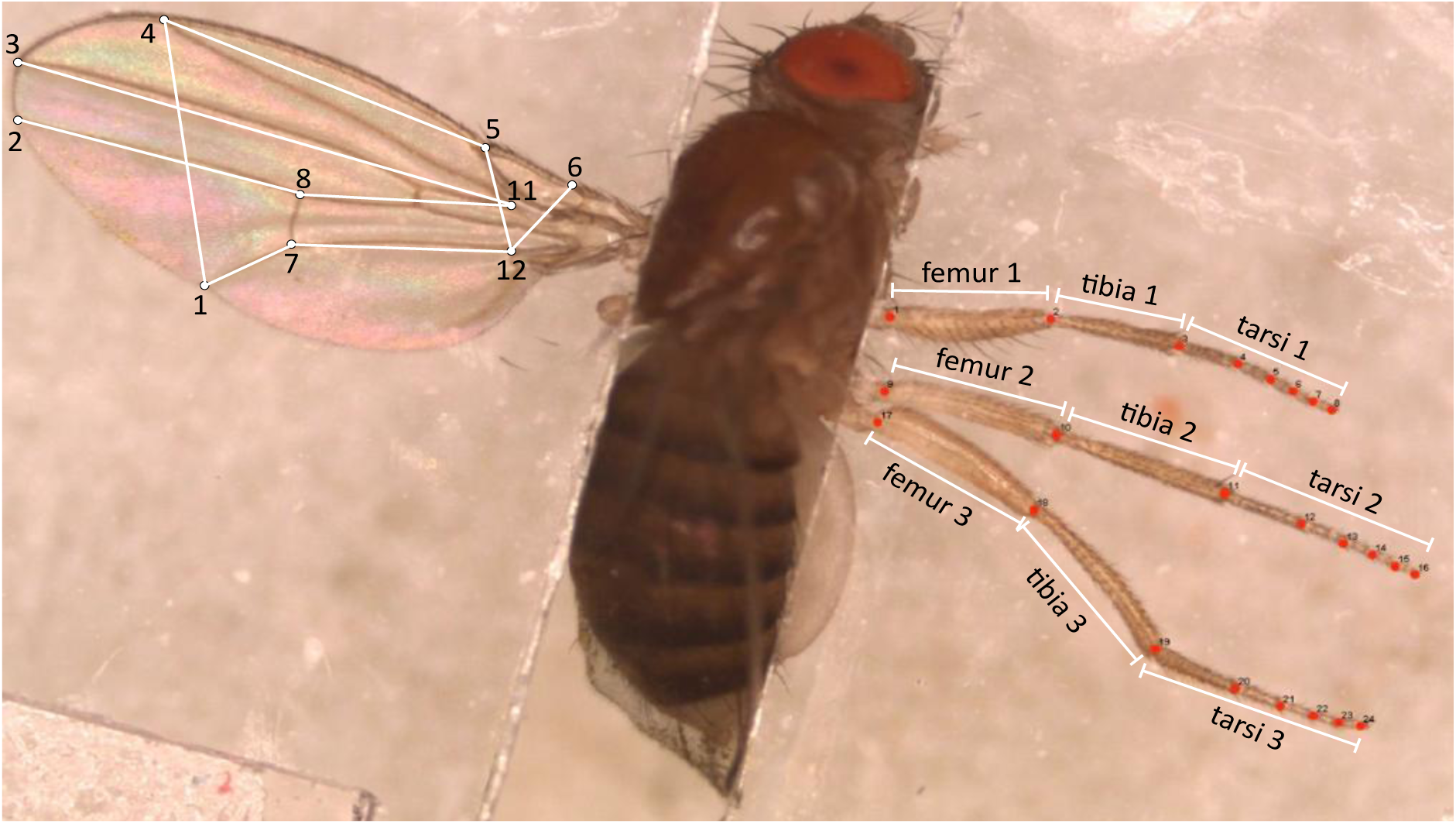
-Anatomical landmarks positioned in the wings and legs of *D. melanogaster* with the distances analyzed.

### 2.4 Leg phenotyping

The machine-learning approach used for leg phenotyping is based on ML-morph [25]. ML-morph is an automated approach for landmark placement using a combination of object detection followed by shape prediction. In this case, due to the high standardization of the fly imaging setup, we skipped the object detection step and directly performed shape prediction using a cascade regression algorithm [27]. The resulting program *appendML* is implemented in python and available on Github repository (https://github.com/agporto/Appendometer).

We trained the algorithm to detect the endpoints of femurs, tibias, and tarsomeres on all three legs (Fig. 3). The machine-learning approach depends on a set of human-marked training images. Training was based on a set of 2,831 marked images that include a variety of Drosophilid fly species, and both high- and low-quality images. A randomly chosen 90% of the available images were used for training (N=2,547), and the remainder were used for validation (N=284). Training images were marked as described above and training was performed using the standard ML-morph pipeline. Given the large number of images in this dataset, we used the validation set to perform an exhaustive grid search of the hyperparameter space of the ML model. We used the best performing model among all grid models as the final one.

Program outputs are TPS, CSV or XML format files containing inferred position of leg joints. We calculated the distances between landmarks along each leg, resulting in seven distances per leg: femur, tibia, and five tarsomeres. The total tarsal lengths are the sum of the tarsomere lengths. The set of nine femur, tibia and tarsi lengths (three for each leg) were then subjected to multivariate outlier detection as described in Supplementary Materials.

We also implemented a Drosophila-specific measurement algorithm based on low-level image processing steps, as described in the Supplementary Material. We refer to this as the “algorithmic” approach in what follows.

### 2.6 Wing phenotyping

We programmed a function to extract the wing region of interest from the full image and recalculate the user-entered wing guide landmark coordinates within that region. Wing measurement then proceeded using the algorithm described in [11]. Splining and error correction was implemented in the Java program *Wings* 4.01 [28]. The program CPR [29] was used to extract fitted models from *Wings* output and performs a Procrustes least-squares superimposition [30] prior to subsequent analyses. Since the leg data consists of nine segment lengths, we chose a representative set of nine lengths between vein intersections on the wing for our principal analyses, shown in Fig. 3.

### 2.7 Repeatability experiment

To quantify the biases and errors arising from the Appendometer, we repeatedly measured 25 individuals of *Drosophila melanogaster* from three different long-term lab populations. Each individual was imaged twice on two different macroscopes, by two different observers, resulting in 200 images in total. Measurements of legs were then estimated from each image using both algorithmic and machine-learning software for legs, and the Wings program for wings. Measurement pipelines for the machine-learning leg data and for wings incorporate the ability to edit the data obtained. The output from these programs was edited by two different observers twice, resulting in 32 estimates of leg and wing variables for each individual. The resulting segment dimensions were analyzed in Proc Mixed in SAS [31].

### 2.9 Species data

We measured wings and legs of 43 Drosophilid taxa, including 17 species in the subgenus Drosophila, 25 species of the subgenus Sophopohora, and an outgroup, *Scaptomyza latifa-siciaeformis*. Supplementary Table S1 provides the sample sizes for each species. We used the [32] phylogeny for all analyses (Fig. 4).

**FIGURE 4.**
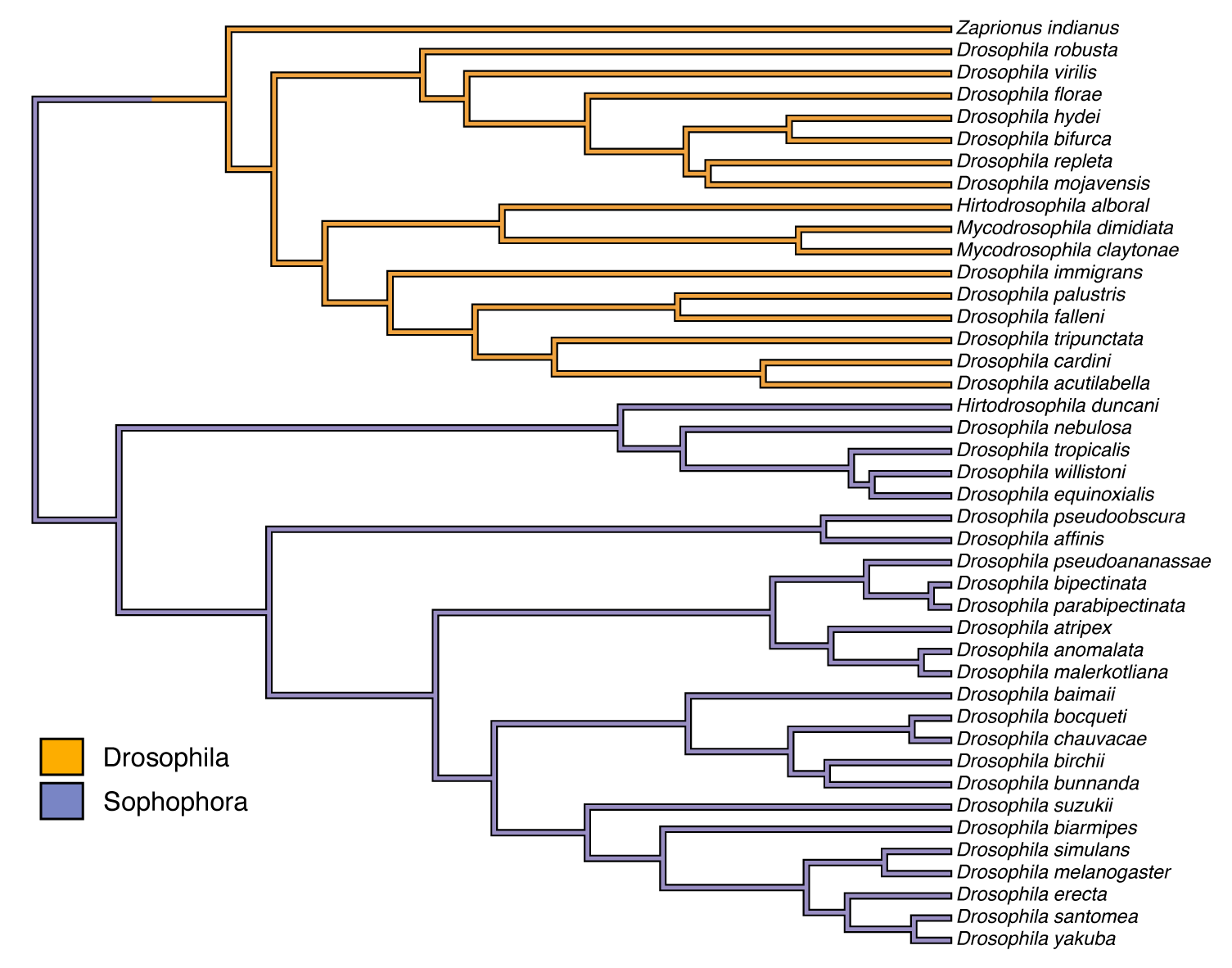
-Phylogeny adapted from [32] to include just the studied species. The outgroup, *Scaptodrosophila latifasciaeformis,* is not included.

### 2.10 Multivariate analyses

We focused our comparative analyses on proportional changes in appendage dimensions by calculating measures of appendage size and shape. Appendage sizes were calculated as the total leg length and the total of the nine wing distances shown in Fig. 3. We then calculated measures of shape as proportions, trait length divided by the appropriate appendage-specific size. Multivariate analyses were carried out on shape, or shape-size, the nine proportions, plus the natural log of total appendage size. The shape data sets are less than full rank, so we subjected them to a principal component analysis, and then scored all specimens on the first eight eigenvectors. To report the results, we rotated the results back to the space of the original variables.

### 2.11 Phylogenetic mixed model

To estimate the rates of evolution in leg and wing morphology across the phylogeny, we used a phylogenetic mixed model [33, 34] under a Brownian motion model of evolution. We estimated rate matrices in Wombat [35] for leg and wing data considering all species together and the subgenus Drosophila and Sophophora separately. We included the sex of the fly as a fixed effect. We compared the fits of full and reduced-rank model likelihoods [36–38], and selected the best-fitting rank model based on Akaike’s Information Criterion corrected for small sample size (AICc).

### 2.12 Within-species variation

We estimated the phenotypic variance/covariance phenotypic matrices (**P**) of leg measurements for the 33 species for which 90 or more individuals were measured. Estimates of **P** were calculated after removing the fixed effect of sex within each taxon. The pooled within-group **P**-matrices for the 35 species with N ≥90 for legs was based on the residuals from a linear model with sex, species and their interaction.

We compared **P**-matrices using random skewers [39, 40]. Each **P**-matrix is multiplied by 10,000 random vectors obtained from a multivariate normal distribution and normalized to a length of one. The average correlation (cosine of the angle) between the response vectors in matrices *i* and *j* to the same skewer is our measure of matrix similarity, *r_ij_*. Sampling error of each matrix puts an upper limit to *r_ij_*. To compensate for this, we bootstrapped the sex-corrected observations for each P-matrix 10,000 times, then calculated the correlation of response vectors for random pairs of these samples. The average of these is the self-correlation, *s_i_*. We calculate the error-adjusted response correlation as 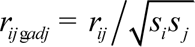.

## 3 RESULTS

### 3.1 Throughput, error and repeatability

We estimated throughput for the combination of wing and leg phenotyping based on two large experiments that utilized many users and together measured over 66,000 flies, as described in Supplementary Results. The entire process of imaging, measurement and error checking for both legs and wings takes an average of 2.7 minutes/per specimen for experienced users. For comparison, hand-landmarking of the 24 leg points shown in Figure 2b with proper attention to accuracy takes from 50 to 60 additional seconds per specimen. Even attentive and careful users occasionally mismark landmarks, resulting in outliers. Hence a hand-marking pipeline properly includes the error correction step. Thus hand-landmarking will add from 0.8 to 1 minute per specimen, decreasing throughput by at least 30%.

The amount of error in human-placed landmarks is a good benchmark for the performance of automated phenotyping. We compared hand-landmarking to automated landmarking for 200 images of legs of three different *Drosophilid* species recorded over a four-year period. Overall, the median distance between leg landmarks hand-marked twice in the same images is 3.0 pixels, averaged over all 24 landmarks. Median distances between replicate landmarks varied from 2.2 pixels for the ends of the terminal tarsomeres to 4.5, 5.0 and 4.1 pixels for the trochanter-femur boundary of the first to third legs respectively. The larger error for this boundary is due to the length of this suture, and the fact that it is nearly perpendicular to the femur axis (Fig. 3 and Supporting Information Video 1). Median differences in segment lengths averaged just 1.4 pixels, a fraction of the point-wise discrepancies, because segment length is relatively insensitive to landmark placement along the segment boundaries.

The median distance between hand-placed and ML-placed landmarks, averaged across all points, was 5.3 pixels, which is 61% greater than the median distance for repeated hand-landmarking. Considering only the 12 landmarks that delineate the ends of the leg segments the error was 5.0 pixels, 48% higher than hand-landmarking. The means and proportional deviations of the estimates of femur, tibia and tarsal lengths are shown in Supplementary Table S2. On average, the means differ by 1.27% from hand-marking.

To investigate the causes of variation in leg measurements, we did a full factorial experiment in which 25 *D. melanogaster* individuals were each imaged twice by two different operators on two different scopes. Results are shown in Table 1. Overall, the among-fly variance was relatively low, with a coefficient of variation (CV) averaging just 3.6% for segment lengths across both measurement approaches. For comparison, within sex CV for leg dimensions in the experiment used to estimate throughput averages 5.1%, suggesting that the among-fly variance in a typical experiment is approximately double that in the repeatability experiment (Table 1). Despite the low among-fly variance, the average repeatability of femur lengths was 75% and averaged 84% for the other segments. Editing of outlier landmark placements greatly improved repeatabilities, from 39% for the raw output of *appendML* to the final average value of 82%.

**Table 1.** Sources of variation in leg dimensions (in *μ*m) using the machine-learning approach.

| Length | Mean | Fixed effects |  |  |  | Variances† |  |  |  | CVs (%) |  |  |
| --- | --- | --- | --- | --- | --- | --- | --- | --- | --- | --- | --- | --- |
|  |  | Sex | Imager | Scope‡ | editor | Fly | image | Error | Repeata- | Fly | image | Error |
|  |  | (F-M) |  |  |  |  |  |  | bility (%) |  |  |  |
| Femur 1 | 536.9 | 7.0 | 3.6* | -4.4** | -0.5 | 372.6 | 87.5 | 50.1 | 73.03 | 3.60 | 1.74 | 1.32 |
| Tibia 1 | 467.5 | 8.7* | -0.8 | 4.5*** | -1.4*** | 237.2 | 43.1 | 29.1 | 76.66 | 3.29 | 1.40 | 1.15 |
| Tarsi 1 | 553.7 | -2.3 | 1.5 | 1.1 | -1.0** | 455.5 | 55.5 | 23.4 | 85.24 | 3.85 | 1.35 | 0.87 |
| Femur 2 | 617.0 | 17.9** | 0.3 | -6.2*** | 1.7*** | 284.8 | 88.1 | 35.1 | 69.80 | 2.74 | 1.52 | 0.96 |
| Tibia 2 | 615.3 | 5.7 | -3.2** | -0.4 | -0.2 | 452.2 | 34.5 | 31.2 | 87.31 | 3.46 | 0.95 | 0.91 |
| Tarsi 2 | 724.6 | 1.0 | 1.5 | -3.4** | -3.0*** | 873.5 | 44.9 | 33.2 | 91.79 | 4.08 | 0.92 | 0.80 |
| Femur 3 | 646.6 | 18.9** | 1.7 | -1.8 | -2.4*** | 430.1 | 57.2 | 41.7 | 81.30 | 3.21 | 1.17 | 1.00 |
| Tibia 3 | 648.2 | 11.9 | -1.2 | -1.8 | 0.0 | 594.2 | 80.3 | 50.6 | 81.95 | 3.76 | 1.38 | 1.10 |
| Tarsi 3 | 798.4 | -5.6 | 1.1 | -5.3** | -1.7** | 1140.6 | 97.7 | 67.1 | 87.38 | 4.23 | 1.24 | 1.03 |

†All variance components significant at P<0.001. ‡ High-resolution Leica -lower-resolution Optem. \* P<0.05; \*\* P<0.01; \*\*\*P<0.0001

Substantial variance is caused by variation among images, with a CV averaging 1.3%. Leg position varies from image to image, including possible stretching of joints, and is likely responsible for this variation. Editors favored minor differences in placement of many landmarks, resulting in an average difference of 0.2% of segment lengths (Table 1). The scope used significantly affected several segment lengths, but these are inconsistent in direction. Sources of variation in leg dimensions using the algorithmic approach are presented in Supplementary Table S3.

Principal component analysis of segment lengths shows 60% of the variance in segment lengths is captured in PC1, suggesting substantial variation in overall size. For most applications of the Appendometer, the shape of legs, variation in relative dimensions, is more interesting than variation in overall size. To quantify error and repeatability in shape, we standardized each segment length by the total lengths of segments. Analyses of these proportions is shown in Supplementary Table S4 for the machine-learning estimates (see Table S5 for algorithmic approach). As expected, leg proportions are quantified less precisely, with average repeatabilities of 40%. The error in estimating femur proportions is particularly high.

For comparison with the leg results, we calculated nine distances among landmarks on wings from the same individual flies. On average, these distances had a repeatability of 87%, somewhat higher than the leg lengths (Supplementary Table S6). Supplementary Table S7 shows repeatabilities of wing distances expressed as a proportion of the total set of distances, analogous to our measure of leg shape. Repeatabilities of proportional distances average 58% over all wing distances, substantially lower than repeatabilities of wing distances and higher than leg proportion repeatabilities.

### 3.3 Rate of evolution

#### 3.3.1 Evolutionary rates of legs and wings

The evolutionary rate matrix for leg size and proportions is shown in Table 2. The rates showed a consistent negative correlation between femur and tibia relative to tarsal proportions, and positive correlations of rates of femur and tibia evolution. This suggests that natural selection tends to favor changes in the relative length of proximal and distal segments of the legs, rather than simple changes in the length of individual segments. At least part of this pattern is due to allometry. Proportional lengths of tibias and femurs were negatively correlated, and tarsal proportions were positively correlated with leg length (Table 2).

**Table 2.**
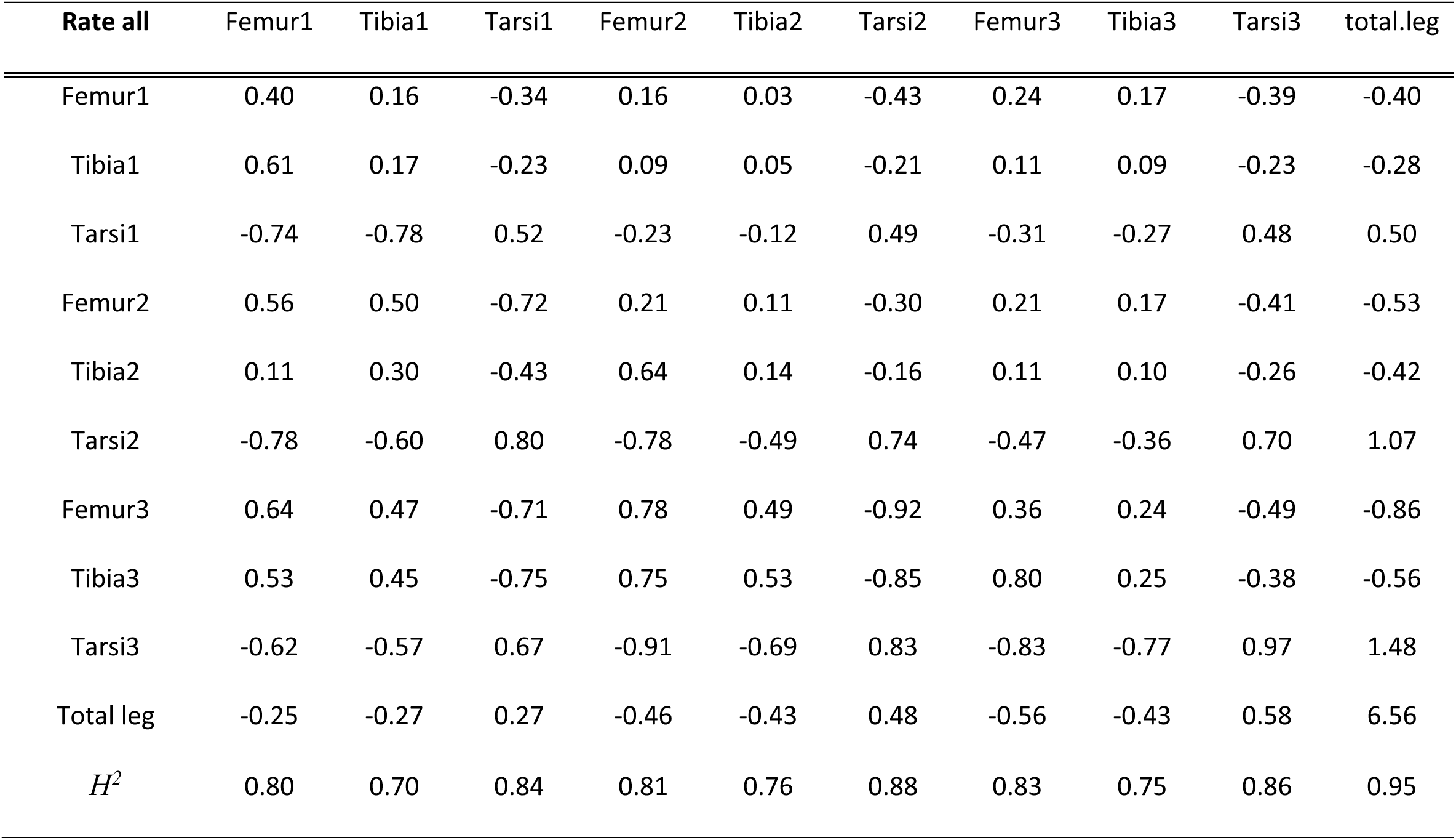
Rates and covariation of rates for back transformed leg data. Diagonal elements repre-sent rates of increase in among species variance for individual traits, while upper off-diagonal elements represent the estimated evolutionary covariance between those traits. Lower off-diago-nal elements are trait correlations. Phylogenetic heritability (*H^2^*) for each trait is in the last row of the table.

| **Rate all** | Femur1 | Tibia1 | Tarsi1 | Femur2 | Tibia2 | Tarsi2 | Femur3 | Tibia3 | Tarsi3 | total.leg |
| --- | --- | --- | --- | --- | --- | --- | --- | --- | --- | --- |
| Femur1 | 0.40 | 0.16 | -0.34 | 0.16 | 0.03 | -0.43 | 0.24 | 0.17 | -0.39 | -0.40 |
| Tibia1 | 0.61 | 0.17 | -0.23 | 0.09 | 0.05 | -0.21 | 0.11 | 0.09 | -0.23 | -0.28 |
| Tarsi1 | -0.74 | -0.78 | 0.52 | -0.23 | -0.12 | 0.49 | -0.31 | -0.27 | 0.48 | 0.50 |
| Femur2 | 0.56 | 0.50 | -0.72 | 0.21 | 0.11 | -0.30 | 0.21 | 0.17 | -0.41 | -0.53 |
| Tibia2 | 0.11 | 0.30 | -0.43 | 0.64 | 0.14 | -0.16 | 0.11 | 0.10 | -0.26 | -0.42 |
| Tarsi2 | -0.78 | -0.60 | 0.80 | -0.78 | -0.49 | 0.74 | -0.47 | -0.36 | 0.70 | 1.07 |
| Femur3 | 0.64 | 0.47 | -0.71 | 0.78 | 0.49 | -0.92 | 0.36 | 0.24 | -0.49 | -0.86 |
| Tibia3 | 0.53 | 0.45 | -0.75 | 0.75 | 0.53 | -0.85 | 0.80 | 0.25 | -0.38 | -0.56 |
| Tarsi3 | -0.62 | -0.57 | 0.67 | -0.91 | -0.69 | 0.83 | -0.83 | -0.77 | 0.97 | 1.48 |
| Total leg | -0.25 | -0.27 | 0.27 | -0.46 | -0.43 | 0.48 | -0.56 | -0.43 | 0.58 | 6.56 |
| $H^2$ | 0.80 | 0.70 | 0.84 | 0.81 | 0.76 | 0.88 | 0.83 | 0.75 | 0.86 | 0.95 |

The evolutionary rate matrix for wing size and proportions is shown in Table 3. The rate of evolution for wing and leg size is very similar, but the estimated rate of wing proportion evolution is higher than for legs. The rates of wing distance proportion evolution are quite variable, suggesting that which distances are analyzed may affect this conclusion. To get a more comprehensive measure of wing rates, we analyzed the rates of evolution for all 66 possible distances between the 12 wing landmarks, measured as proportions of the total of the 66 distances with results shown in Supplementary Table S8. The mean rate of evolution over all proportions was 0.089, (S.D. 0.050), while the comparable rates for leg proportions were lower and much less variable (0.055, S.D. 0.016). The source of this variation is the strong allometry of wing shape [17, 18], where points 1, 4, 7 and 8 move distally relative to the remainder of the points as size increases (see Figure 3 for anatomical landmarks visualization). This results in higher evolutionary rates for distances between these wing points and other points.

**Table 3.** Rate of evolution and traits correlation of back transformed wing data including all spe-cies.

| Rate all | w1_4 | w1_7 | w2_8 | w3_11 | w4_5 | w5_12 | w6_12 | w7_12 | w8_11 | total.wing |
| --- | --- | --- | --- | --- | --- | --- | --- | --- | --- | --- |
| w1_4 | 0.40 | 0.01 | -0.05 | -0.12 | -0.16 | 0.07 | 0.14 | -0.17 | -0.11 | -0.59 |
| w1_7 | 0.02 | 0.82 | 1.00 | 0.20 | -0.43 | -0.05 | 0.03 | -0.75 | -0.83 | -0.87 |
| w2_8 | -0.06 | 0.79 | 1.93 | 0.53 | -0.91 | -0.11 | -0.07 | -1.02 | -1.30 | -1.69 |
| w3_11 | -0.33 | 0.38 | 0.67 | 0.33 | -0.46 | -0.05 | -0.05 | -0.20 | -0.18 | -0.43 |
| w4_5 | -0.21 | -0.38 | -0.54 | -0.65 | 1.49 | -0.14 | -0.13 | 0.36 | 0.38 | 1.48 |
| w5_12 | 0.31 | -0.17 | -0.24 | -0.25 | -0.33 | 0.12 | 0.05 | 0.06 | 0.05 | 0.12 |
| w6_12 | 0.72 | 0.09 | -0.16 | -0.30 | -0.35 | 0.50 | 0.10 | -0.06 | -0.01 | -0.15 |
| w7_12 | -0.28 | -0.86 | -0.77 | -0.37 | 0.31 | 0.19 | -0.19 | 0.92 | 0.86 | 1.00 |
| w8_11 | -0.17 | -0.86 | -0.88 | -0.30 | 0.29 | 0.15 | -0.02 | 0.85 | 1.13 | 1.13 |
| Total wing | -0.37 | -0.38 | -0.49 | -0.30 | 0.48 | 0.14 | -0.19 | 0.42 | 0.42 | 6.30 |
| $H^2$ | 0.89 | 0.94 | 0.96 | 0.91 | 0.94 | 0.75 | 0.84 | 0.93 | 0.94 | 0.77 |

#### 3.3.2 Evolutionary rates in Drosophilid subgenera

Independent analyses of each Drosophilid subgenera revealed that Sophophora presented a higher rate of evolution than Drosophila for the leg data (Table 4). For the wing data, Sophophora and Drosophila showed similar rates of evolution (Table 4).

**Table 4.** Trace and average phylogenetic heritability for proportions and proportions + size.

| | Trace Pro-<br>portions | Trace Prop.+Size | Mean Phylo. $H^2$<br>Prop | Mean Phylo. $H^2$<br>Prop+size |
| --- | --- | --- | --- | --- |
| Legs all | 3.76 | 10.3 | 0.8 | 0.82 |
| Wings all | 2 | 8.33 | 0.86 | 0.85 |
| Legs Drosophila | 2.28 | 12.5 | 0.7 | 0.72 |
| Legs Sophophora | 4.86 | 9.21 | 0.85 | 0.86 |
| Wings Drosophila | 2.18 | 13.23 | 0.83 | 0.83 |
| Wings Sophophora | 2.13 | 6.53 | 0.88 | 0.86 |

Phylogenetic heritability (*H^2^*, Table 4) was slightly higher for wing proportions (mean 0.86) compared to leg proportions (mean 0.80). Analyses for the leg data of each subgenus revealed that Sophophora had a substantially higher *H^2^*than Drosophila (0.86 vs. 0.70 for proportions, Table 4). For the wing data, *H^2^* was again higher in Sophophora than in Drosophila but the difference was much smaller (0.88 vs. 0.88 for proportions, Table 4).

### 3.4 Covariation between legs and wings

One of the key questions that the Appendometer allows us to investigate is whether the rates of wing and leg evolution are correlated. Table 5 shows the correlations of leg and wing trait evolution. Leg and wing size are relatively highly correlated, and both measures of size have substantial correlations with proportions in the other appendage. The cross-appendage correlations show intriguing correspondences with the negative relationships between tarsal vs. tibia and femur proportions seen in the leg data. Tarsal proportions are positively correlated with total wing size, while femur and tibia proportions are negatively correlated with wing size (Table 5). This pattern of correlation is repeated for proportional distances along the length of the wing (strongly for more distal lengths w2_8, w3_11, w4_5, and weakly for the proximal lengths w7_12 and w8_11). Three measures of wing width, w1_4, w6_12 and w5_12, show the converse pattern of positive correlations with femur and tibia and negative with tarsal proportions (Table 5). To test how much of this is due to allometry, we reestimated the wing/leg correlations for residuals from regressions of proportions on the log of the sum of wing and leg size. As shown in Supplementary Table S9, the pattern of size-adjusted correlations was on average similar in intensity to those in Table 4 (R^2^ Table 4=0.079, R^2^ Table S9=0.065), although in many cases the nature of the correlations with size removed is rather different.

**Table 5.** Correlations of the rate of leg and wing trait evolution.

|  | w1_4 | w1_7 | W2_8 | W3_11 | W4_5 | W5_12 | W6_12 | W7_12 | w8_11 | total.wing |
| --- | --- | --- | --- | --- | --- | --- | --- | --- | --- | --- |
| Femur1 | 0.33 | -0.13 | -0.30 | -0.36 | -0.12 | 0.28 | 0.59 | 0.09 | 0.12 | -0.27 |
| Tibia1 | 0.56 | -0.15 | -0.22 | -0.44 | -0.19 | 0.19 | 0.30 | 0.05 | 0.06 | -0.52 |
| Tarsi1 | -0.45 | 0.10 | 0.26 | 0.44 | 0.12 | -0.33 | -0.54 | 0.01 | -0.06 | 0.37 |
| Femur2 | 0.46 | 0.09 | -0.01 | -0.29 | -0.28 | 0.38 | 0.52 | -0.18 | -0.18 | -0.40 |
| Tibia2 | 0.26 | 0.19 | 0.13 | -0.04 | -0.26 | 0.19 | 0.06 | -0.19 | -0.20 | -0.26 |
| Tarsi2 | -0.32 | -0.15 | 0.09 | 0.26 | 0.22 | -0.33 | -0.61 | 0.14 | 0.12 | 0.28 |
| Femur3 | 0.27 | 0.32 | 0.10 | -0.17 | -0.27 | 0.22 | 0.60 | -0.27 | -0.29 | -0.23 |
| Tibia3 | 0.20 | 0.18 | -0.06 | -0.16 | -0.15 | 0.24 | 0.47 | -0.18 | -0.09 | -0.17 |
| Tarsi3 | -0.41 | -0.19 | -0.08 | 0.19 | 0.34 | -0.23 | -0.45 | 0.20 | 0.22 | 0.38 |
| total.leg | -0.51 | -0.46 | -0.44 | -0.02 | 0.46 | 0.14 | -0.16 | 0.48 | 0.56 | 0.54 |

To investigate the degree to which selection on one appendage can be expected to influence the other appendage, we compared the sizes of the sub-matrices of the rate covariance matrix capturing leg, wing or leg-wing covariances, using the Frobenius norm as a measure of matrix size [41]. As noted above for the matrix traces, the wing sub-matrix is larger than the leg sub-matrix. We can, however, compare the magnitude of leg-wing covariances to those of each appendage to get a rough idea of how important correlated responses would be. For proportional distances, the leg and wing sub-matrices are both larger than the off-diagonal leg-wing matrix, by 2.1 times (C.I. 1.0-3.0) legs, and 6.8 times (C.I. 3.5-8.7) for wings, suggesting the correlated responses will be fairly modest. Size-corrected proportional distances yield similar results (Supplementary Table S9). On the other hand, for shape plus size, the leg sub-matrix is about equal to the size of the off-diagonal leg-wing matrix (ratio 1.085, C.I. 1.8-3.3), while wing sub-matrix is 2.6 times the size of the off-diagonal leg-wing matrix (C.I. 1.9-4.1). Thus, selection on size of appendages will generate large, correlated responses on the other appendage relative to selection on appendage shape. For other sets of proportional wing distances, these ratios can be rather different. The bottom line is that correlated responses of one appendage to selection on the other can be substantial for some combinations of traits.

### 3.5 Within-species variation, and the relationships between P- and R-matrices

We compared species **P**-matrices, the pooled **P**-matrix and the **R**-matrix using random skewers, with the results shown in Appendix 1, Tables A1 for legs, A2 for wings. All matrices were accurately estimated with a median self-similarity of 0.97 for wing and leg matrices, and 0.99 for rate matrices. The self-similarity corrected skewer correlations showed that P-matrices were substantially similar for both legs (median 0.87, IQ range 0.12) and wings (median 0.88, IQ range 0.12).

The rate matrix was on average similar to the P-matrices (median skewer correlation for legs 0.66, IQ range 0.14, median for wings 0.79, IQ range 0.10), slightly less similar than P-matrices are to each other. Figure 5 shows that there is a strong relationship between the log variance component in the R-matrix and the log variance component in the pooled P-matrix for both legs (b=1.51; p=0.001; R^2^=0.78) and wings (b=1.90; p=0.00; R^2^=0.75).

**FIGURE 5.**
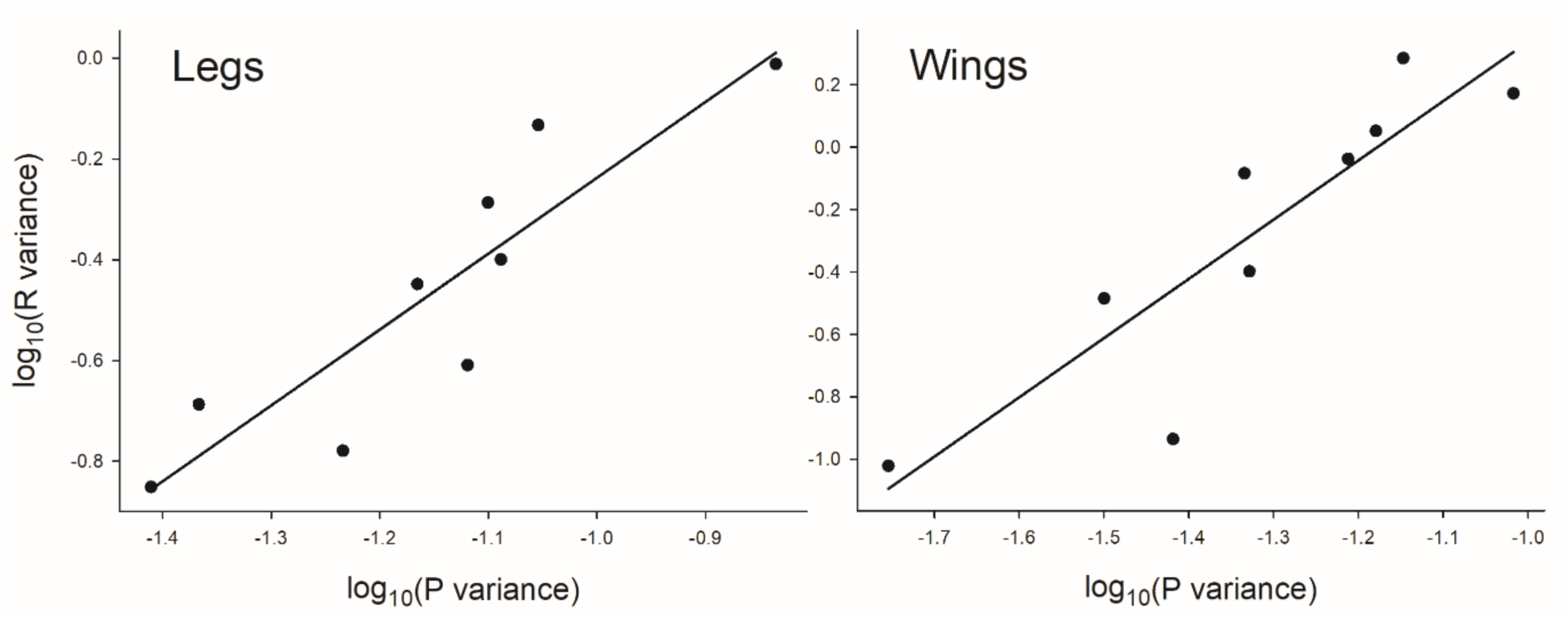
– Relationship between the log variance component in the rate matrix and the log variance component in the pooled P-matrix for legs and wings.

## 4 DISCUSSION

We have developed a system that enables rapid, repeatable measurement of legs and wings of Drosophilid flies, and use it to measure the patterns of within- and among-species variation in these developmentally related appendages. We begin by discussing the capabilities and limitations of the measurement system, followed by an examination of the variation and evolution of legs and wings.

The hardware used for imaging is simple to construct, relatively inexpensive, and is readily adaptable to other winged insect taxa. The machine-learning image processing software is generally applicable to any 2-D images that capture the locations of morphological features. It has already been applied to a wide range of traits in diverse taxa [25].

The key hardware is a pair of suction devices that simultaneously immobilize and present one wing and one set of legs for simultaneous imaging. Image analysis software enables rapid measurements of wings and legs to be obtained. For experienced users, the average time to obtain and check high-dimensional data on both wings and legs is less than three minutes per specimen, two minutes to obtain an image, and then an average of about a minute inspecting and correcting measurements on outlier specimens. The recovered data includes the lengths of femurs, tibia and tarsomeres on all three legs on one side of a fly, and a detailed model of wing vein placement. Wing measurement is accomplished using previously developed software [11, 28].

We successfully trained the machine-learning algorithm on data from many flies in the sub-family Drosophilinae. We were then able to use this approach on a wide sample of species from that sub-family. Legs in this sub-family are similar enough that the algorithm seemed equally successful when applied to each species (data not shown). In addition, we have used the Appendometer system in two very large, but as-yet-unpublished experiments in *Drosophila melanogaster* and *D. simulans*, for which we measured wings and legs for over 66,000 flies.

Our results do show that the machine-learning algorithm increases the variance of repeat leg measurements based on new images of about 60% over hand-landmarking of leg segments on the same images. This increase in error needs to be balanced against the 30% decrease in throughput, labor costs for the landmarking process, and dealing with user differences in mean landmark positions in large experiments. Our use of the machine learning algorithm rather than hand-landmarking enabled us to complete a large selection experiment (Jones and Houle unpublished) that required imaging and measurement of over 1,000 flies per week for 8 months. Hand-landmarking would have added 17 hours per week to this process, a burden that would have required hiring an additional worker or decreasing the number of flies measured.

Our experiment to estimate the repeatability of measurements from the Appendometer showed that the size was quite accurately measured for both legs and wings. On the other hand, the repeatability for measures of shape, form holding size constant, was substantially lower, averaging a modest 40%. However, the sample of flies used in this experiment was unusually uniform in size and shape, only half the true among-individual variation that we estimated for an outbred lab population. For a sample of flies with among individual variance similar to this population, repeatability would be expected to rise to 55%.

Before adopting the flexible machine-learning approach as our favored method for leg measurement, we developed an algorithmic approach that combines low-level image processing with a priori information on the expected leg morphology to trace the length and widths of the leg segments. This method is described in the Supplementary Material. We ultimately abandoned the algorithmic approach as it did not return reasonable results for five to six percent of specimens, which then must be treated as missing data. In addition, extension of that algorithm to different taxonomic groups would require additional programming, and not just retraining of working code. The algorithmic approach does have several advantages. It eliminates observer effects on the recovered data, and repeatably estimates the widths of femurs and tibias. An im-proved algorithm could potentially outperform the machine-learning approach.

### Variation and evolution of fly legs

To demonstrate the power of the Appendometer, we analyzed the patterns of within species variation and evolution of legs in the sub-family *Drosophilinae* in the family *Drosophilidae*. We obtained data for 43 species. Within taxon variation revealed a striking pattern where the proportion of each leg in the tarsal segments was negatively correlated with the proportion of the leg made up of femur and tibia. Femur and tibia proportions were always positively correlated. In addition, these patterns were also prominent for segments across all three legs: for example, tarsal proportions for each leg were negatively correlated with the femur and tibia of all three legs. Homologous leg segments were always positively correlated.

To study the pattern of evolutionary changes in leg proportions, we estimated the rates of evolution of segment proportions and leg size using a phylogenetic mixed model. These rates revealed the same pattern of correlations observed within species: The rate of evolution of tarsal proportions was negatively correlated with the femur and tibia proportions, suggesting an evolutionary tradeoff where changes in one necessitated compensatory change in the other. This is partly driven by allometry, as larger species showed that tarsi make up an increased proportion of leg length in species with larger body size. This sort of pattern is to some extent necessitated by quantifying shape as a proportion of total leg length, as increases in one proportion require decreases in other proportions. Under a simple model of the evolution of proportions, increases in any one proportion would be somewhat negatively correlated with the evolution of all other proportions. However, this is not observed between tibia and femur proportions, which are strongly positively correlated. This suggests that tarsal proportions are more negatively correlated with tibia and femur proportions that predicted under a simple null model.

In addition, we used the combined wing and leg data to describe the patterns of variational and evolutionary covariation between wings and legs. As noted in the Introduction, one could expect either that wing and leg evolution might be highly correlated due to the extensive sharing of the developmental pathways and genes expressed during their development [22–24], or largely independent, as the function and presumably natural selection on each appendage are not directly related.

Our results show real but modest correlations between wing and leg evolution. One factor driving evolutionary correlations between wings and legs is allometric changes in appendage shape [17, 18]. The sizes of wings and legs are positively correlated, and the size of each appendage is also modestly correlated with the proportions of features in the other appendage. On the other hand, when we removed allometry from the matrices using size as a predictor variable in our analysis of evolutionary rates, the ratios of the total sizes of the leg- and wing-specific co-variance matrices to the between leg- and wing-covariances matrices was substantially unchanged. Evolutionary correlations remain even after allometry is removed. While the cross-appendage correlations were modest in size, it does suggest that either natural selection or the pattern of within species covariation does modestly entangle the evolution of the two appendages.

Houle et al. [18] has previously found that the amount of mutational and genetic variation for wing shape in *Drosophila melanogaster* predicts the pattern of evolution within the family *Drosophilidae*, a pattern repeated within sepsid flies [42]. More generally, **P** matrices for other *Drosophilid* species are highly correlated with genetic and mutational variation in *D. melanogaster* (Tsuboi et al., manuscript in preparation). We found that the same pattern holds between phenotypic variation and evolutionary rate for leg shape in our the more limited sample of taxa measured here. Similar patterns have been observed in other taxa [43–46], suggesting that the pattern of correlations between variation and evolution is quite general. The explanation for this pattern is unknown, as it can be explained either when variation constrains the rate of evolution, or by reshaping of the pattern of variation to correspond to evolutionary rates [46, 47].

## DECLARATIONS

### Ethics approval and consent to participate

Not applicable

### Consent for publication

Not applicable

### Availability of data and materials

Data, R scripts, and the program *appendML* are available on Github repository: (https://github.com/agporto/Appendometer). Additional supporting information can be found in the online version of this article.

### Authors’ contributions

DH conceived the project.

CSM measured wing and leg morphology of the species data set.

DMR and DH edited the leg and wing data sets.

DMR and CSM conducted the comparative analyses of P matrices and evolutionary rates.

DMR designed and carried out the repeatability experiment.

DMR and DH analyzed the repeatability data.

AP implemented the ML algorithm for legs.

DR and DH wrote the manuscript.

All authors contributed critically to the drafts and gave final approval for publication.

## Acknowledgements

This work was supported by grants from the National Science Foundation (DEB-1556774, to D.H) and a NVIDIA Hardware Grant to A.P. Thanks to Bill Green for writing the algorithmic leg phenotyping software, and to Luke Jones, Dominick Trozzo-Stamper, Ryan Fortune, Brad Salemi, and Kevin Doheny for helping to run the various projects that use the Appendometer, and the 20 FSU undergraduates who each measured some of the flies analyzed here. We also thank Claudia Russo and Beatriz Mello for providing phylogenetic information on Drosophilid flies, and Dean Adams for discussions on evolutionary rate estimates using mixed models and PGLS.

