## Supplementary Material for "Appendometer: A system for simultaneous, high-throughput morphometry of *Drosophila* legs and wings"

*Methods in Ecology and Evolution*

### 1. Supplementary Methods

#### *1.1 Specimen Handling*

Maneuvering the left wing and right set of legs into the wing grabbers, as shown in Fig. 2a and b in the main text, is a delicate process that requires some practice to master. The process begins with the wing grabber in position on the left-hand end of the channel on the positioning block over the surface of a $CO_2$ diffuser (Flystuff Flypad, Genesee Scientific), which is used to hold the specimen at the same level as the grabbers. The imager uses a paintbrush to move the anaesthetized specimen close to the grabber, and then extend the left wing until the suction of the wing grabber pulls it into the imaging slot. A leg grabber is then placed in the right-hand slot in the positioning block and slid to the left until it approaches the right-hand set of legs of the immobilized fly. The imager uses the paint brush to coax the legs into an extended position and into the grabber, which is then moved further to the left until the femur, tibia, and tarsi of all three legs are within the grabber. The diffuser is then withdrawn, and the positioning block with the immobilized fly in the grabbers is moved to the imaging area on the left side of the plexiglass sheet shown in Fig. 2c, where the user takes an image.

We intended this system to be used with active $CO_2$ anesthesia, but positioning of individual flies in the grabbers can take several minutes. This can result in morbidity and mortality. Consequently, we usually use FlyNap (Carolina Biological Supply Company) for anesthesia. FlyNap anaesthetized flies stay anaesthetized for up to 30 minutes without apparent damaging side effects. This makes it possible to simultaneously immobilize small groups of flies, reducing fly handling time.

While the fly handling, imaging and data recording tasks are readily accomplished by a single user, the rate of imaging can be increased if one user positions flies in the grabbers, while the other records specimen information and digitizes the guide landmarks.

#### *1.2 Algorithmic leg phenotyping*

Algorithmic phenotyping is implemented in the Java program Legs, written by Bill Green. Program outputs are a file containing inferred position of leg elements, and an image overlaying some of these on the portion of the region of interest contain the legs (Fig. 2b). Output information includes the pixels closest to the central leg axes, inferred positions of leg joints and ends, smoothed mean, maximum and minimum widths, and the arc length from the femur landmarks to the femur/tibia joint, from the femur/tibia joint to the proximal end of the first tarsomere, and the sum of the arc length along the tarsomeres to the terminal claw on each leg.

The algorithm proceeds in the following major steps:

1. The locations of the user-placed landmarks on the trochanter-femur boundary of each leg are used to extract a region of interest containing the legs and excluding the rest of the image.

2. The region of interest is thresholded to identify possible leg elements. There is chromatic aberration in the images unless focus is precise over the entire image. The program first chooses which channel to apply the algorithm to. The blue color channel is most informative when there is little visual contrast within each leg; the red channel is most informative when there is substantial contrast between leg edges and center. The chosen color channel is thresholded using Otsu's method, then holes are filled. Dark pixels above the threshold are the foreground pixels that may lie within the legs.

3. To identify the central axis of each leg, and to estimates leg widths, boundary pixels in the foreground are used to construct a Voronoi diagram using Fortune's algorithm. The foreground is then skeletonized using inverse medial axis transformation. This skeleton is then progressively pruned of short elements. Once pruning is complete the central axis is inferred from those remaining edges whose boundary points are on opposite sides of the foreground region – the putative sides of the leg segment. The inferred widths and center-axis pixels have local variation due to foreground edge properties caused by small features such as foreground hairs or extraneous material on the image. To smooth out these noisy features the central axis is approximated by a C1 cubic spline.

4. The proximal regions of the femurs frequently touch, so their widths can be confounded. To allow the proximal center axis of femurs to be approximated, a vector between the basal landmark of that femur and the estimated center axis out to a fixed distance away from that landmark is substituted for the skeletonized axis.

5. The joints between femur and tibia, and between tibia and the proximal tarsal segment as local minima of the ratio of the smoothed width derivative relative to the smoothed width. Search for these minima was conducted only at proportions of the total leg length within the

range of human-validated joint locations. When several minima existed within the region examined, the program chooses the one with the highest value of the smoothed direction derivative, indicating a flexed joint. The above algorithms work well when the interior angle of orientation between leg segments is greater than 90 degrees. When the angles are less than 90 degrees, the inferred widths of the segment tips near the joint are distorted. Cases with larger angles were identified by unusually large local changes in the smoothed width. The joint was then inferred to fall at the point where the orthogonal direction to the central axis spline becomes atypical of the rest of that segment.

6. Program parameters specify the minimum and maximum widths of leg segments, allowing detection of regions that either cannot be legs, or capture portions of image containing extraneous elements such as dirt or hairs in the image. Non-leg regions are discarded based on their length-width ratios, and contiguity to other putative leg segments working outwards from the proximal landmark locations. Once this process is complete, cubic splines for the central leg axis are re-estimated. When the boundary has a gap due to pruning, the width is estimated by interpolation of neighboring non-missing boundaries. The smoothed width at each point on the splined axis is then estimated from the intersection of the vector orthogonal to the axis with the boundaries of the foreground.

7. Segment lengths are then estimated as the arc length between joints along the splined central axis fit to each leg.

#### *1.3 Error detection and correction*

For all data sets, we used robust outlier detection routines to rank specimens by their degree of unusualness. For the leg segments, outliers were detected using the function covMcd in the R package robustbase. The images overlaid with the fitted model are then shown to the user in reverse order of their Mahalanobis distance from the robust mean, which are then discarded or corrected as described below. Errors were corrected until the majority of specimens require minimal correction, usually achieved after examining 10-20% of the images thereby minimizing the number of images the user examines. Hand-correction of point placement for specimens at typical Mahalanobis distances does not lead to gains in repeatability because intra-observer error in point placement is magnified. Using this approach, the time needed for error checking for a set of flies is far less than the aggregate imaging time. Images of specimens with missing body parts, or that are out of focus or incorrectly exposed are always discarded.

#### *1.4 Algorithmic leg phenotyping*

In a few percent of images, *Legs* fails to fit a model that falls within the range of plausible leg morphologies, in which case Legs does not return a model. This can sometimes be remedied by repositioning the guide landmarks, but in other cases there are no obvious flaws that can explain the lack of fit. Errors in guide landmark placement were corrected in tpsDig2 (Rohlf 2017), and *Legs* was then rerun, and outlier detection repeated. The editor sorts the outlier images into two categories 1) incorrect placement of leg joint landmarks by the algorithm, and 2) biological outliers with unusual segment lengths. Images in category 1 were discarded; biological outliers were retained for further analysis.

#### *1.5 Machine-learning leg phenotypes*

In the machine-learning pipeline, the recovered data are 24 landmark locations, so we implemented presentation and error-correction in a digitization program, tpsDig2 (Rohlf 2017), and data can be obtained for all specimens with good images.

#### *1.6 Wing phenotypes*

Outlier detection and robust covariance matrix estimation was carried out for each sex on the full set of landmarks, and on the wing outline only, using the MVE algorithm (Van Der Linde 2004), implemented in *Wings* 4.01. *Wings* 4.01 enables the user to correct the splines by dragging the spline control points, or re-spline the image with different processing parameters to obtain a good model fit. The program *CPR* (Márquez 2012–2014) was used to extract fitted models from Wings output, derive the positions of landmarks and semi-landmarks from the splines, perform Procrustes superimposition, slide the semi landmarks, and to perform additional outlier detection using an ordination of the data.

### 2. Supplementary Results

#### *2.1 Algorithmic phenotyping*

The error classification step typically takes 10 minutes for a set of 500 flies, and there is no opportunity for error correction. The Legs program failed to return a model for 2.6% of usable images, and 3.0% of the fitted models did not correctly reflect leg morphology.

The 12 Alg landmarks averaged 5.2 pixels distance from the corresponding hand-landmarked ones, slightly less accurate than the ML approach. Our analysis of the Alg lengths and widths from the repeatability experiment is shown in Table S3. The single residual error term estimated for the Alg data includes both the effects of the image and any error introduced by the algorithm. Average repeatability was 74% for femur lengths, and 90.3% for the remaining segment lengths. This suggests that landmark placements were slightly more consistent using the Alg approach, except for the femur-trochanter boundary where user-placed guide landmarks are treated as data, in contrast to the ML data where the femur-trochanter landmark is included in the training set. Consistent with this, the total of the two ML error components, image, and residual error, was slightly higher than the single Alg error component. This suggests that the error in ML measurements is increased by the error correction step, relative to the Alg results. Imager had large effects on femur lengths in the Alg data. Repeatabilities of femur and tibia widths average 73%, but just 38% for tarsi. The lower resolution images had substantially higher estimated widths.

Analyses of leg proportions is shown in Table S5 for the Alg data. Leg proportions are quantified less precisely than lengths, with average repeatabilities of 55% in the Alg data, and 31% for femur proportions. These repeatabilities are superior to the average of 33% achieved by the ML algorithm.

### *2.2 Throughput*

We estimated throughput for the Appendometer from two large experiments that employed numerous part-time undergraduate assistants. In the first such experiment, we imaged 30,393 individuals for wings and legs in a *Drosophila melanogaster* population over 85 weeks for a quantitative genetic and genome-wide association experiment (Houle, unpublished). In the second, we performed two simultaneous artificial selection experiments in *D. melanogaster* and its sister species *D. simulans*, for which we imaged a total of 36,255 flies (Jones and Houle, unpublished). The average imaging time was 2.2 minutes for single imagers and 1.7 minutes when one user manipulated the flies, while the other placed the guide landmarks and recorded specimen information. After the initial 10 weeks of the quantitative genetic experiment, a total of 1.8% of the images were unusable because the image did not include all three legs or focus or exposure was inadequate. In the selection experiment just 0.5% were unusable.

Experience using the Appendometer was the most important determinant of imaging rate; three imagers who recorded less than 25 images averaged 6.2 minutes/fly, while three users that recorded more than 4000 images averaged 2.0 minutes/fly. The mean rate of imaging by 16 users in the data set was a linear function of the $\log_{10}$ number of total images recorded by that imager ($b$=-1.60, $P$<0.0001, $R^2$=0.88). The time required to fit the leg model for sets of 500 images is less than a minute.

The entire process of phenotyping and error checking for both wings and legs can be accomplished in an average of 2.7 minutes/specimen by experienced users.

**Table S1.** Drosophilid species and their respective subgenera classification. Total number of individuals measured for legs (N total legs) and wings (N total wings) and number of individuals by sex (**N♀♀**: female; **N♂♂**: male).

| Species | N total legs | N♀♀ legs | N♂♂ legs | N total wings | N♀♀ wings | N♂♂ wings | Subgenera |
|---|---|---|---|---|---|---|---|
| *Drosophila acutilabella* | 118 | 59 | 59 | 118 | 59 | 59 | Drosophila |
| *Drosophila affinis* | 102 | 44 | 58 | 199 | 99 | 100 | Sophophora |
| *Drosophila anomalata** | 117 | 57 | 60 | 117 | 58 | 59 | Sophophora |
| *Drosophila atripex* | 100 | 49 | 51 | 98 | 47 | 51 | Sophophora |
| *Drosophila baimaii* | 17 | 9 | 8 | 16 | 9 | 7 | Sophophora |
| *Drosophila biarmipes* | 118 | 59 | 59 | 120 | 60 | 60 | Sophophora |
| *Drosophila bifurca* | 100 | 45 | 55 | 101 | 45 | 56 | Drosophila |
| *Drosophila bipectinata* | 99 | 49 | 50 | 85 | 47 | 38 | Sophophora |
| *Drosophila birchii* | 100 | 50 | 50 | 97 | 47 | 50 | Sophophora |
| *Drosophila bocqueti* | 99 | 50 | 49 | 95 | 49 | 46 | Sophophora |
| *Drosophila bunnanda** | 97 | 47 | 50 | 90 | 46 | 44 | Sophophora |
| *Drosophila cardini* | 116 | 58 | 58 | 119 | 59 | 60 | Drosophila |
| *Drosophila chauvacae* | 100 | 50 | 50 | 100 | 50 | 50 | Sophophora |
| *Drosophila equinoxialis* | 90 | 46 | 44 | 98 | 48 | 50 | Sophophora |
| *Drosophila erecta* | 100 | 50 | 50 | 98 | 50 | 48 | Sophophora |
| *Drosophila falleni** | 122 | 62 | 60 | 122 | 62 | 60 | Drosophila |
| *Drosophila florae** | 13 | 5 | 8 | 13 | 5 | 8 | Drosophila |
| *Drosophila hydei* | 191 | 92 | 99 | 199 | 100 | 99 | Drosophila |
| *Drosophila immigrans* | 194 | 99 | 95 | 198 | 99 | 99 | Drosophila |
| *Drosophila malerkotliana** | 100 | 50 | 50 | 92 | 49 | 43 | Sophophora |
| *Drosophila melanogaster* | 130 | 65 | 65 | 130 | 65 | 65 | Sophophora |
| *Drosophila mojavensis* | 99 | 54 | 45 | 99 | 54 | 45 | Drosophila |
| *Drosophila nebulosa* | 99 | 49 | 50 | 100 | 50 | 50 | Sophophora |
| *Drosophila palustris* | 84 | 45 | 39 | 84 | 45 | 39 | Drosophila |
| *Drosophila parabipectinata* | 99 | 50 | 49 | 98 | 50 | 48 | Sophophora |
| *Drosophila pseudoananassae* | 100 | 50 | 50 | 91 | 48 | 43 | Sophophora |
| *Drosophila pseudoobscura* | 100 | 50 | 50 | 100 | 50 | 50 | Sophophora |
| *Drosophila repleta* | 4 | 1 | 3 | 4 | 1 | 3 | Drosophila |
| *Drosophila robusta* | 118 | 60 | 58 | 120 | 60 | 60 | Drosophila |

| | | | | | | | |
|---|---|---|---|---|---|---|---|
| *Drosophila santomea* | 101 | 53 | 48 | 102 | 53 | 49 | Sophophora |
| *Drosophila simulans* | 196 | 100 | 96 | 197 | 98 | 99 | Sophophora |
| *Drosophila suzukii** | 196 | 96 | 100 | 192 | 100 | 92 | Sophophora |
| *Drosophila tripunctata* | 117 | 58 | 59 | 120 | 60 | 60 | Drosophila |
| *Drosophila tropicalis* | 100 | 59 | 41 | 100 | 59 | 41 | Sophophora |
| *Drosophila virilis* | 100 | 50 | 50 | 100 | 50 | 50 | Drosophila |
| *Drosophila willistoni* | 93 | 45 | 48 | 93 | 45 | 48 | Sophophora |
| *Drosophila yakuba* | 100 | 50 | 50 | 98 | 49 | 49 | Sophophora |
| *Hirtodrosophila alboral* | 27 | 8 | 19 | 27 | 8 | 19 | Drosophila |
| *Hirtodrosophila duncani** | 2 | 2 | 0 | 2 | 2 | 0 | Sophophora |
| *Mycodrosophila claytonae** | 3 | 2 | 1 | 3 | 2 | 1 | Drosophila |
| *Mycodrosophila dimidiata** | 25 | 8 | 17 | 25 | 8 | 17 | Drosophila |
| *Scaptodrosophila latifasciaeformis* | 126 | 70 | 56 | 124 | 69 | 55 | Scaptodrosophila |
| *Zaprionus indianus* | 120 | 60 | 60 | 72 | 35 | 37 | Drosophila |
| **Sum (Σ)** | 4232 | 2115 | 2117 | 4256 | 2149 | 2107 | |

*Species not presented in Russo's phylogeny Russo et al (2013). Sister species were used to substitute the species in the phylogeny.

**Table S2.** Discrepancies in leg segment lengths derived from hand vs. algorithmic methods.

| | mean length (px) | | | Difference in mean lengths (%) | | Median error (% of hand-estimate) | |
|---|---|---|---|---|---|---|---|
| | Hand | Alg | ML | Alg/hand | ML/hand | Alg/hand | ML/hand |
| Front femur | 188.6 | 189.6 | 191.7 | 0.53 | 1.66 | 1.73 | 2.13 |
| Front tibia | 165.2 | 164.6 | 163.9 | -0.36 | -0.77 | 1.84 | 2.00 |
| Front tarsi | 195.0 | 203.7 | 194.3 | 4.47 | -0.35 | 4.52 | 1.62 |
| Middle femur | 217.1 | 221.0 | 221.1 | 1.79 | 1.83 | 2.19 | 2.35 |
| Middle tibia | 226.9 | 224.8 | 222.6 | -0.94 | -1.90 | 1.41 | 1.86 |
| Middle tarsi | 241.8 | 248.5 | 244.2 | 2.76 | 0.99 | 2.70 | 1.45 |
| Rear femur | 228.5 | 229.2 | 229.5 | 0.28 | 0.43 | 1.75 | 1.72 |
| Rear tibia | 232.7 | 234.5 | 228.8 | 0.78 | -1.66 | 1.44 | 2.01 |
| Rear tarsi | 255.9 | 264.5 | 260.7 | 3.36 | 1.88 | 3.16 | 2.14 |

**Table S3.** Sources of variation in leg dimensions (in μm) using the algorithmic approach.

| | | | Fixed effects | | | Variances† | | | CV (%) | |
|---|---|---|---|---|---|---|---|---|---|---|
| L or W‡ | Segment | Mean | Sex (F-M) | Imager | Scope‡ | Fly | Error | Repeat-ability (%) | Fly | Error |
| L | Femur 1 | 539.5 | 17.2* | 12.4*** | 0.2 | 413.6 | 154.0 | 72.9 | 3.77 | 2.30 |
| L | Tibia 1 | 467.6 | -3 | 2.0* | -1.9 | 220.7 | 42.3 | 83.9 | 3.18 | 1.39 |
| L | Tarsi 1 | 584.8 | 1.7 | 2.2* | 5.8** | 412.6 | 50.0 | 89.2 | 3.47 | 1.21 |
| L | Femur 2 | 625.6 | 15.9* | 10.8*** | 1.5 | 344.1 | 152.2 | 69.3 | 2.97 | 1.97 |
| L | Tibia 2 | 624.7 | 4.6 | -2.9** | -2.3* | 464.3 | 37.0 | 92.6 | 3.45 | 0.97 |
| L | Tarsi 2 | 741.0 | -6.3 | 0.8 | 5.9*** | 831.9 | 75.4 | 91.7 | 3.89 | 1.17 |
| L | Femur 3 | 646.4 | 19.9** | 2.2 | -0.2 | 473.0 | 131.1 | 78.3 | 3.36 | 1.77 |
| L | Tibia 3 | 668.8 | 9.2 | 0.8 | -0.8 | 569.1 | 53.5 | 91.4 | 3.57 | 1.09 |
| L | Tarsi 3 | 822.2 | -22.3** | 0.6 | 4.5** | 1294.7 | 95.8 | 93.1 | 4.38 | 1.19 |
| W | Femur 1 | 40.0 | -0.18 | 0.44* | -1.92*** | 5.70 | 1.80 | 76.0 | 5.97 | 3.35 |
| W | Tibia 1 | 24.4 | -0.28 | -0.01 | -1.67*** | 2.32 | 0.56 | 80.6 | 6.24 | 3.07 |
| W | Tarsi 1 | 15.3 | -0.65** | 0.38** | -1.75*** | 0.24 | 0.64 | 27.3 | 3.20 | 5.23 |
| W | Femur 2 | 36.7 | 0.36 | 0.50** | -1.07*** | 2.9 | 1.4 | 67.6 | 4.62 | 3.20 |
| W | Tibia 2 | 25.1 | 0.12 | 0.05 | -1.52*** | 2.2 | 0.7 | 75.4 | 5.88 | 3.36 |
| W | Tarsi 2 | 14.6 | 0.42 | 0.59*** | -1.69*** | 0.4 | 0.7 | 35.8 | 4.22 | 5.65 |
| W | Femur 3 | 39.5 | 1.32 | 0.96** | -1.05*** | 4.1 | 3.0 | 57.9 | 5.13 | 4.38 |
| W | Tibia 3 | 27.4 | 0.74 | 0.30* | -1.67*** | 2.2 | 0.9 | 70.5 | 5.38 | 3.48 |
| W | Tarsi 3 | 17.6 | 0.43 | 0.25 | -1.79*** | 0.7 | 0.7 | 48.9 | 4.72 | 4.82 |

†All variance components significant at $P<0.001$.

‡ L=segment length; W=mean segment width

§ High-resolution Leica -lower-resolution Optem

**Table S4.** Sources of variation in leg segment proportions using the machine-learning approach.

| Segment | Mean % | Variances† Fly | image | Error | Rep. (%) | Fly CV (%) | Image CV (%) | Error CV (%) |
|---|---|---|---|---|---|---|---|---|
| Femur 1 | 9.59 | 0.0211 | 0.0374 | 0.0166 | 28.1 | 1.5 | 2.0 | 1.3 |
| Tibia 1 | 8.34 | 0.0088 | 0.0202 | 0.0118 | 21.6 | 1.1 | 1.7 | 2.1 |
| Tarsi 1 | 9.87 | 0.0533 | 0.0152 | 0.0071 | 70.5 | 2.3 | 1.2 | 1.5 |
| Femur 2 | 11.01 | 0.0169 | 0.0254 | 0.0108 | 31.9 | 1.2 | 1.4 | 1.7 |
| Tibia 2 | 10.98 | 0.0232 | 0.0182 | 0.0125 | 43.0 | 1.4 | 1.2 | 1.6 |
| Tarsi 2 | 12.94 | 0.0726 | 0.0381 | 0.0113 | 59.5 | 2.1 | 1.5 | 1.7 |
| Femur 3 | 11.54 | 0.0225 | 0.0251 | 0.0132 | 37.0 | 1.3 | 1.4 | 1.7 |
| Tibia 3 | 11.57 | 0.0259 | 0.0365 | 0.0196 | 31.6 | 1.4 | 1.7 | 2.0 |
| Tarsi 3 | 14.25 | 0.0515 | 0.0540 | 0.0276 | 38.7 | 1.6 | 1.6 | 2.0 |

†All variance components significant at $P<0.001$.

**Table S5.** Sources of variation in leg segment proportions using the algorithmic approach.

| | | Variances† | | | CV (%) | |
|---|---|---|---|---|---|---|
| Segment | Mean % | Fly | Error | Rep. (%) | Fly | Error |
| Femur 1 | 9.43 | 0.0207 | 0.0464 | 30.8 | 1.53 | 2.28 |
| Tibia 1 | 8.17 | 0.0073 | 0.0121 | 37.6 | 1.05 | 1.35 |
| Tarsi 1 | 10.22 | 0.0492 | 0.0143 | 77.5 | 2.17 | 1.17 |
| Femur 2 | 10.94 | 0.0187 | 0.0359 | 34.2 | 1.25 | 1.73 |
| Tibia 2 | 10.92 | 0.0324 | 0.0113 | 74.1 | 1.65 | 0.97 |
| Tarsi 2 | 12.95 | 0.0745 | 0.0207 | 78.3 | 2.11 | 1.11 |
| Femur 3 | 11.30 | 0.0137 | 0.0338 | 28.8 | 1.04 | 1.63 |
| Tibia 3 | 11.69 | 0.0170 | 0.0132 | 56.3 | 1.12 | 0.98 |
| Tarsi 3 | 14.37 | 0.0787 | 0.0272 | 74.3 | 1.95 | 1.15 |

†All variance components significant at $P<0.001$.

**Table S6.** Sources of variation in distances between wing landmarks.

| Length | Mean | Fixed effects | | | | Variances† | | | Repeata-bility (%) | CVs (%) | | |
|---|---|---|---|---|---|---|---|---|---|---|---|---|
| | | Sex (F-M) | Imager | Scope‡ | editor | Fly | image | Error | | Fly | image | Error |
| d1_4 | 972.7 | 9.6 | -2.1 | -21.1*** | 6.1*** | 3269.1 | 59.7 | 109.0 | 95.09 | 5.88 | 0.79 | 1.07 |
| d1_7 | 324.2 | 1.2 | -1.4 | -7.0*** | 6.7*** | 535.0 | 29.8 | 98.9 | 80.61 | 7.13 | 1.68 | 3.07 |
| d2_8 | 939 | 9.0 | -1.1 | -14.3*** | 3.3*** | 3154.9 | 37.6 | 19.5 | 98.22 | 5.98 | 0.65 | 0.47 |
| d3_11 | 1698.6 | 16.1 | -2.4 | -2.4 | 6.9*** | 9513.6 | 209.8 | 165.2 | 96.21 | 5.74 | 0.85 | 0.76 |
| d4_5 | 1167.5 | -7.4 | -2.5 | -22.7*** | 17.0*** | 7257.9 | 73.4 | 555.5 | 92.03 | 7.30 | 0.73 | 2.02 |
| d5_12 | 368.9 | 26.2*** | -0.2 | -6.9*** | 4.4*** | 360.9 | 68.2 | 61.9 | 73.50 | 5.15 | 2.24 | 2.13 |
| d6_12 | 281.6 | 19.6*** | 1.2 | -3.8** | -2.0*** | 234.5 | 58.9 | 24.3 | 73.81 | 5.44 | 2.73 | 1.75 |
| d7_12 | 719.5 | 7.8 | -2.8 | -0.9 | 4.3*** | 2180.2 | 103.8 | 93.2 | 91.71 | 6.49 | 1.42 | 1.34 |
| d8_11 | 713.5 | 18.4* | -1.0 | 15.6*** | 6.5*** | 1660.5 | 133.1 | 137.4 | 85.99 | 5.71 | 1.62 | 1.64 |

†All variance components significant at P<0.001.
‡ High-resolution Leica -lower-resolution Optem.
* P<0.05; ** P<0.01; ***P<0.0001

**Table S7.** Sources of variation in distances between wing landmarks as a proportion of the sum of all landmark distances.

| | | Variances† | | | | CVs (%) | | |
|---|---|---|---|---|---|---|---|---|
| Length | Mean (%) | Fly | image | Error | Repeata-bility (%) | Fly | image | Error |
| pd1_4 | 13.54 | 0.0748 | 0.0083 | 0.0151 | 76.14 | 2.02 | 0.67 | 0.91 |
| d1_7 | 4.51 | 0.0540 | 0.0050 | 0.0086 | 79.97 | 5.15 | 1.56 | 2.05 |
| d2_8 | 13.07 | 0.0783 | 0.0050 | 0.0088 | 85.02 | 2.14 | 0.54 | 0.72 |
| d3_11 | 23.64 | 0.0220 | 0.0091 | 0.0207 | 42.53 | 0.63 | 0.40 | 0.61 |
| d4_5 | 16.23 | 0.0451 | 0.0175 | 0.0716 | 33.62 | 1.31 | 0.81 | 1.65 |
| d5_12 | 5.13 | 0.0144 | 0.0119 | 0.0121 | 37.53 | 2.34 | 2.12 | 2.15 |
| d6_12 | 3.91 | 0.0101 | 0.0112 | 0.0062 | 36.73 | 2.57 | 2.70 | 2.02 |
| d7_12 | 10.01 | 0.0845 | 0.0156 | 0.0164 | 72.52 | 2.90 | 1.25 | 1.28 |
| d8_11 | 9.93 | 0.0543 | 0.0152 | 0.0189 | 61.38 | 2.35 | 1.24 | 1.38 |

**Table S8.** Mean-standardized rates of evolution of 66 distances on the wing. Distances were standardized first by dividing by the total of the 66 distances, and then to a mean value of 1.0. Points identified in Fig. 3.

| Point 1 | Point 2 | Rate | S.E. |
|---|---|---|---|
| 1 | 2 | 0.073 | 0.012 |
| 1 | 3 | 0.057 | 0.010 |
| 1 | 4 | 0.049 | 0.009 |
| 1 | 5 | 0.031 | 0.006 |
| 1 | 6 | 0.026 | 0.006 |
| 1 | 7 | 0.225 | 0.022 |
| 1 | 8 | 0.116 | 0.014 |
| 1 | 9 | 0.064 | 0.010 |
| 1 | 10 | 0.052 | 0.009 |
| 1 | 11 | 0.030 | 0.006 |
| 1 | 12 | 0.036 | 0.007 |
| 2 | 3 | 0.171 | 0.014 |
| 2 | 4 | 0.181 | 0.025 |
| 2 | 5 | 0.033 | 0.007 |
| 2 | 6 | 0.024 | 0.006 |
| 2 | 7 | 0.093 | 0.017 |
| 2 | 8 | 0.112 | 0.019 |
| 2 | 9 | 0.056 | 0.012 |
| 2 | 10 | 0.053 | 0.011 |
| 2 | 11 | 0.028 | 0.007 |
| 2 | 12 | 0.030 | 0.007 |
| 3 | 4 | 0.253 | 0.032 |
| 3 | 5 | 0.031 | 0.007 |
| 3 | 6 | 0.021 | 0.005 |
| 3 | 7 | 0.077 | 0.014 |
| 3 | 8 | 0.099 | 0.018 |
| 3 | 9 | 0.049 | 0.011 |
| 3 | 10 | 0.048 | 0.010 |
| 3 | 11 | 0.025 | 0.006 |
| 3 | 12 | 0.025 | 0.006 |
| 4 | 5 | 0.080 | 0.016 |
| 4 | 6 | 0.062 | 0.014 |
| 4 | 7 | 0.051 | 0.009 |
| 4 | 8 | 0.074 | 0.012 |
| 4 | 9 | 0.069 | 0.013 |
| 4 | 10 | 0.074 | 0.014 |
| 4 | 11 | 0.067 | 0.015 |
| 4 | 12 | 0.059 | 0.013 |

| Point 1 | Point 2 | Rate | S.E. |
|---|---|---|---|
| 5 | 6 | 0.075 | 0.008 |
| 5 | 7 | 0.083 | 0.013 |
| 5 | 8 | 0.094 | 0.014 |
| 5 | 9 | 0.107 | 0.010 |
| 5 | 10 | 0.133 | 0.011 |
| 5 | 11 | 0.099 | 0.009 |
| 5 | 12 | 0.065 | 0.007 |
| 6 | 7 | 0.069 | 0.013 |
| 6 | 8 | 0.076 | 0.014 |
| 6 | 9 | 0.096 | 0.013 |
| 6 | 10 | 0.098 | 0.012 |
| 6 | 11 | 0.083 | 0.007 |
| 6 | 12 | 0.080 | 0.008 |
| 7 | 8 | 0.147 | 0.012 |
| 7 | 9 | 0.154 | 0.020 |
| 7 | 10 | 0.131 | 0.018 |
| 7 | 11 | 0.086 | 0.014 |
| 7 | 12 | 0.088 | 0.014 |
| 8 | 9 | 0.158 | 0.019 |
| 8 | 10 | 0.146 | 0.018 |
| 8 | 11 | 0.098 | 0.016 |
| 8 | 12 | 0.091 | 0.015 |
| 9 | 10 | 0.184 | 0.009 |
| 9 | 11 | 0.172 | 0.018 |
| 9 | 12 | 0.134 | 0.015 |
| 10 | 11 | 0.170 | 0.018 |
| 10 | 12 | 0.118 | 0.013 |
| 11 | 12 | 0.109 | 0.007 |

**Table S9.** Correlations of evolutionary rate for appendage proportions once allometry is removed.

| | w1_4 | w1_7 | W2_8 | W3_11 | W4_5 | W5_12 | W6_12 | W7_12 | w8_11 |
|---|---|---|---|---|---|---|---|---|---|
| Femur1 | 0.19 | -0.24 | -0.39 | -0.36 | 0.12 | 0.31 | 0.56 | 0.25 | 0.31 |
| Tibia1 | 0.34 | -0.45 | -0.52 | -0.46 | 0.27 | 0.30 | 0.36 | 0.45 | 0.44 |
| Tarsi1 | -0.21 | 0.21 | 0.37 | 0.38 | -0.18 | -0.36 | -0.57 | -0.17 | -0.26 |
| Femur2 | -0.11 | 0.01 | -0.09 | -0.18 | 0.02 | 0.34 | 0.45 | -0.01 | -0.02 |
| Tibia2 | -0.23 | 0.15 | 0.08 | 0.07 | -0.10 | 0.17 | -0.01 | -0.10 | -0.10 |
| Tarsi2 | 0.14 | -0.02 | 0.19 | 0.24 | -0.04 | -0.31 | -0.47 | -0.05 | -0.09 |
| Femur3 | -0.29 | 0.23 | 0.05 | -0.09 | -0.07 | 0.16 | 0.45 | -0.15 | -0.15 |
| Tibia3 | -0.31 | 0.16 | -0.05 | -0.08 | 0.01 | 0.16 | 0.32 | -0.12 | -0.02 |
| Tarsi3 | 0.25 | -0.08 | 0.04 | 0.10 | 0.04 | -0.21 | -0.33 | 0.01 | 0.03 |
